# *Cryptococcus deuterogattii* VGIIa infection associated with travel to the Pacific Northwest outbreak region in an anti-GM-CSF autoantibody positive patient in the United States

**DOI:** 10.1101/493239

**Authors:** Shelly Applen Clancey, Emily J. Ciccone, Marco A. Coelho, Joie Davis, Li Ding, Renee Betancourt, Samuel Glaubiger, Yueh Lee, Steven M. Holland, Peter Gilligan, Julia Sung, Joseph Heitman

## Abstract

The Pacific Northwest (PNW), Vancouver Island, Oregon, and Washington have been the location of an ongoing *Cryptococcus gattii* outbreak since the 1990s, and there is evidence that the outbreak is expanding along the West Coast into California. Here we report a clinical case of a 69-year-old, HIV-negative man from North Carolina who was diagnosed with a fungal brain mass by magnetic resonance imaging (MRI) and pathology. He had traveled to Seattle and Vancouver three years earlier and Costa Rica four months prior to presentation. Phenotypic evidence shows the fungal mass isolated from the patient’s brain is *C. gattii*. In agreement with the phenotypic results, MLST provides genotypic evidence that assigns the infecting organism within in the *C. gattii* species complex and belonging to the *C. deuterogattii* VGIIa clade. Whole genome sequencing revealed >99.99% identity to the *C. deuterogattii* reference strain R265, indicating that the infecting strain is derived from the highly clonal outbreak strains in the PNW. We conclude the patient acquired the *C. gattii* infection during his travel to region three years prior and the infection was dormant for an extended period of time before causing disease. The patient tested positive for anti-granulocyte-macrophage colony-stimulating factor (GM-CSF) autoantibodies, supporting earlier reports that implicate these autoantibodies as a risk factor a risk factor associated with *C. gattii* infection.

**Importance:** Mortality rates associated with *C. gattii* infections are estimated to be between 13% and 33% depending on an individual’s predisposition, and *C. gattii* has caused more at least 39 deaths in the PNW region. There have been four other international travel cases reported in patients from Europe and Asia with travel history to the PNW, but this study describes the first North American travel who acquired *C. deuterogattii* infection presenting within the United States, and the first case of a *C. deuterogattii* outbreak infection associated with anti-GM-CSF autoantibodies. Early and accurate diagnoses are important for disease prevention, treatment, and control of infectious diseases. Continual reporting of *C. deuterogattii* infections is necessary to raise awareness of the ongoing outbreak in the PNW and alert travelers and physicians to the endemic areas with potential risks.

## Introduction

Infectious diseases (e.g. tuberculosis and pneumonia) were the leading cause of death 150 years ago, but at the turn of the 21^st^ century the main cause of death changed from infectious diseases to physiopathological illnesses (e.g. heart disease, stroke, cancer) (1). Despite modern medical advances and successful efforts to reduce the spread of infectious agents, infectious diseases still account for an estimated 13 million deaths per year, compared to the approximately 40 million non-infectious deaths, the majority of which occur in developing countries (2, 3). Globalization and population expansion have increased the spread of infectious diseases despite strong efforts to counter these threats to public health. Additionally, international travel exposes tourists to a variety of diseases and infectious agents they may not otherwise encounter, and with antibiotic and antifungal resistance in human pathogens on the rise, the ability to combat infectious diseases is compromised (4, 5). With the increased mobilization of people, animals, and goods, early recognition among health care providers and the public is an essential component in preventing additional infectious disease cases.

*Cryptococcus* is a fungal pathogen responsible for nearly 220,000 human infections per year (6-8). Among the more than 30 *Cryptococcus* species, *C. neoformans* and *C. gattii* are responsible for the vast majority of human and animal infections. Unlike its sister species, *C. neoformans*, which is dispersed worldwide and infects primarily immunocompromised individuals, certain *C. gattii* subtypes often infect otherwise healthy subjects and are thought to have more restricted geographical niches (9). A 2014 study associated the presence of anti-granulocyte-macrophage colony-stimulating factor (GM-CSF) autoantibodies as a risk factor for central nervous system infections of *C. gattii* but not *C. neoformans*, suggesting that previously unrecognized immune system defects may contribute to disease susceptibility (10). Both *C. neoformans* and *C. gattii* establish infection when desiccated yeast cells or spores are inhaled from the environment into the lungs, in some cases causing pneumonia, and then disseminating to the central nervous system to cause meningoencephalitis. The time from exposure of *C. gattii* to disease is thought to range from 2 to 12 months based on the reported cases of exposure, but may be longer (11).

Historically, all *C. gattii* isolates have been classified as the same species but were separated into five different VG/AFLP molecular types: VGI, VGII, VGIII, VGIV, and AFLP10. New phylogenetic analyses and other evidence supports that each of the VG types actually represent a different species: *C. gattii* (VGI), *C. deuterogattii* (VGII), *C. bacillisporus* (VGIII), *C. tetragattii* (VGIV), and *C. decagattii* (AFLP10) (12, 13). Using multi-locus sequence typing (MLST), each lineage can be further divided into additional molecular subtypes. For example, multiple molecular subtypes exist in *C. deuterogattii*: VGIIa, VGIIb, and VGIIc. Notably, the *MAT***a** allele is underrepresented in all molecular types of the *C. gattii* species complex; the mating type locus (*MAT*) is a gene cluster containing alleles of one of two idiomorphs, *MAT***a** or *MAT*α, and determines mating compatibility (14). *C. gattii* and *C. deuterogattii* have an apparent α unequal number of *MAT***a** and *MAT*α isolates, and very few *MAT***a** strains of these species have been isolated from nature or patients (15). In contrast, the *MAT***a** allele is more highly represented in the VGIIIc subtype of *C. bacillisporus* in that 39% of characterized strains are of the *MAT***a** mating type (16–18). Is it important to characterize each lineage as its own species as it has been shown there is variability in ecological and host niches, virulence, and antifungal susceptibility, resulting in variable success in clinical treatment between the lineages (12).

*C. gattii* has long been a recognized pathogen in South America and Australia, and more recently an outbreak has been recognized on Vancouver Island, Canada, and the United States Pacific Northwest (PNW) causing more than 300 reported human cases and at least 39 deaths, according to the Centers for Disease Control and Prevention (19). Within the last decade, numerous publications have reported genomic and phylogenetic analyses of the outbreak strains, advancing our understanding of the epidemiology and clinical associations of *C. gattii* (20, 21). *C. deuterogattii* has been identified as the causative agent for the ongoing epidemic in the PNW, and continual human and veterinary cases confirm the outbreak is spreading geographically down the West Coast (22, 23). Not only is *C. deuterogattii* a threat to native inhabitants in the PNW and the surrounding areas (e.g. Washington and Oregon), but individuals who travel to the area are also at risk. In fact, travel-related cases have been reported since the outbreak began (24–28), but these appear to be international tourists from Europe and Asia visiting the PNW.

Here we present a case of travel-related cryptococcal central nervous system (CNS) disease in which an individual residing in North Carolina acquired a *C. deuterogattii* infection during travel to the PNW but did not begin to show symptoms until several years after returning home. Unique to this case is the extended incubation period of *C. deuterogattii,* which was three times longer than previously predicted latency periods, before causing disease. Additionally, the patient tested positive for anti-GM-CSF autoantibodies, a suggested risk factor for *C. gattii* infections (10, 29). Although there have been reports of international travel to the PNW resulting in *C. deuterogattii* infections (25, 26, 28), to our knowledge this is the first reported case of a travel-associated cryptococcal CNS disease caused by *C. deuterogattii* from the Pacific Northwest presenting in the United States. Globalization continues to influence the emergence and resurgence of infectious diseases, and continual reporting of travel-related cases of *C. deuterogattii* is crucial to raise awareness of the ongoing fungal outbreak in the PNW and potential risks to travelers. Additionally, providers treating domestic patients should be aware of their travel history, even remote travel years prior to presentation.

## Results

### Clinical presentation and treatment

The patient is a 69-year-old man who presented with three weeks of clumsiness, dizziness, shuffling gait, memory trouble, headache, and worsening fine motor skills. He reported a cough and shortness of breath for 1 to 2 months that had resolved 3 months prior to presentation, and had just returned from an intense bicycling trip that had lasted several days. Because of the new and progressive neurologic symptoms, magnetic resonance imaging (MRI) of the brain with contrast was performed and showed multiple ring-enhancing lesions with associated edema and central necrosis (Figure 1A). The largest lesion was in the right parietal lobe (3.7 × 3.1 cm; Figure 1A, white arrow) with a 3 mm leftward midline shift. Other abnormalities were present in the left inferior temporal lobe (2.2 × 1.8 cm), frontoparietal lobe (1.7 × 1.7 cm), and parietal lobe (2.3 × 2.0 cm), as well as along the midline dura (2.4 × 1.0 cm).

**Figure 1.**
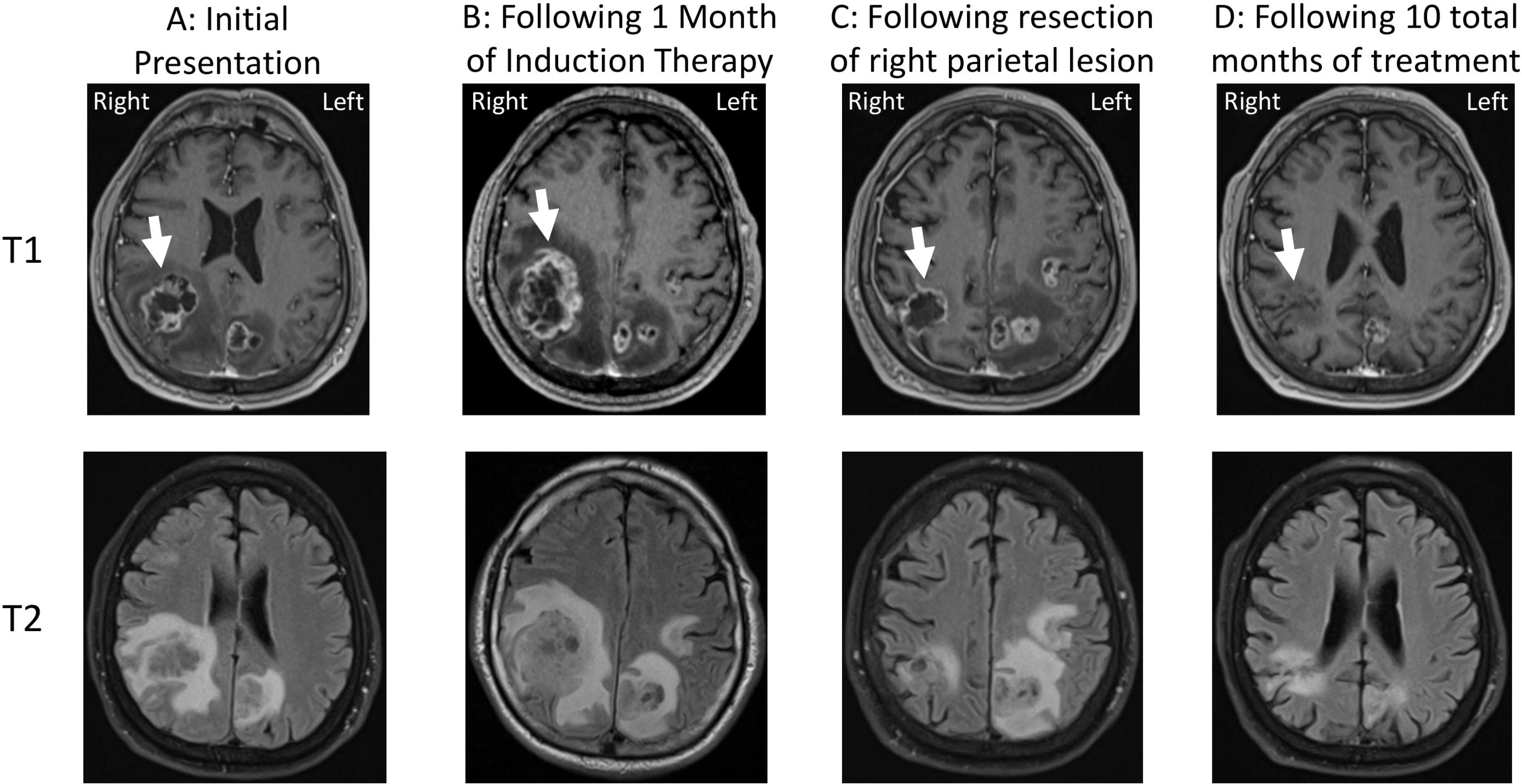
Changes in parietal lobe lesions on brain MRI over the course of treatment. Top panels represent post-gadolinium T1-weighted images and bottom panels represent post-gadolinium T2-weighted images. A) MRI at initial presentation. Circumferential ring enhancement on the post gadolinium T1-weighted axial image and increased signal surrounding the lesion on the axial T2-FLAIR image representing marked vasogenic edema (arrow). The heterogeneously increased signal within the cavity of the lesions represents necrosis. B) MRI following 1 month of induction therapy. Increased T2 signal within the lesion cavity on the FLAIR sequence as well as increased surrounding edema and overall size of the right parietal lesion indicating interval worsening with likely progressive necrosis of brain tissue (arrow). A leftward midline shift was appreciated. C) MRI following resection of the right parietal lesion. Note improvement in the edema surrounding the lesion on the FLAIR sequence and an interval decrease in the size of the right parietal lesion (arrow). D) Subsequent MRI imaging after 10 months total of treatment. Improved T2/FLAIR signal abnormality with decreasing size of the resection cavity as well as decreased size of the other visible lesions (arrow) was observed.

Given concern that the lesions represented metastases, computed tomography of the chest, abdomen, and pelvis was performed, revealing a 2 cm, spiculated right lower lobe nodule consistent in appearance with small cell lung cancer. The lung lesion was not amenable to sampling, so the left temporal lobe mass from the brain was partially resected for definitive diagnosis. Intra-operatively, a frozen section of the firm, grey-colored mass revealed fungal organisms (Figure 2B). Following a review of the patient’s travel history that revealed a trip to Costa Rica four months prior to symptom onset where he visited a bird sanctuary, as well as a trip to Vancouver and Seattle three years prior lasting several weeks, patient serum was sent for cryptococcal antigen testing and returned positive with a titer of 1:1280 (Table 1). The patient was started on 4 mg/kg daily of liposomal amphotericin B (AmB) combined with 2,000 mg four times daily of 5-flucytosine (5-FC) as induction therapy for a presumed cryptococcoma. Additionally, an extended dexamethasone taper was prescribed for cerebral edema (4 mg twice daily for 6 days, then dose halved every 3 days for a total of an 18-day course).

**Table 1.** Patient serum and cerebrospinal fluid (CSF) laboratory results at the time of diagnosis and 4 and 8 weeks post-induction therapy. AST = aspartate aminotransferase, ALT = alanine transaminase, WBC = white blood cell count, ANC = absolute neutrophil count, ALC = absolute lymphocyte count.

|  | *Test*<br>*(normal range)* | *Normal*<br>*range* | *Time of*<br>*diagnosis* | *After 4 weeks*<br>*of induction*<br>*therapy* | *After 8 weeks*<br>*of induction*<br>*therapy* |
| --- | --- | --- | --- | --- | --- |
| Serum | Cryptococcal Antigen Titer | Negative | 1:1280 | 1:320 | 1:2560 |
|  | Creatinine | 0.7-1.30 mg/dL | 0.72 | 1.73 | 1.59 |
|  | AST | 19-55 U/L | 33 | 59 | 27 |
|  | ALT | 19-72 U/L | 31 | 123 | <8 |
|  | WBC | 4.5-11 x10<sup>9</sup> cells/L | 13.3 | 3.9 | 4.5 |
|  | Platelets | 150-440 x10<sup>9</sup> cells/L | 254 | 140 | 174 |
|  | ANC | 2.0-7.5 x10<sup>9</sup> cells/L | 12.2 | 1.9 | 2.2 |
|  | ALC | 1.5-5.0 x10<sup>9</sup> cells/L | 0.6 | 1.3 | 1.7 |
| | Absolute CD4 Count (%) | 510-2,230 cells/μL<br>(34-58%) | 228* (38) | 871 (56) | - |
| | Absolute CD8 Count (%) | 180-1,520 cells/μL<br>(18-38%) | 132* (22) | 358 (23) | - |
| CSF | Opening Pressure | - | 27mmHg | ND |  |
|  | Cryptococcal Antigen | Negative | 1:40 |  |  |
|  | Culture | Negative | Negative |  |  |
| | Nucleated cells | 0-5/μl | 5 | | |
|  | Protein | 15-45 mg/dL | 48 mg/dL |  |  |
|  | Glucose | 50-75 mg/dL | 75 mg/dL |  |  |

\*Testing done three days after initiation of dexamethasone, ND=Not Determined

**Figure 2.**
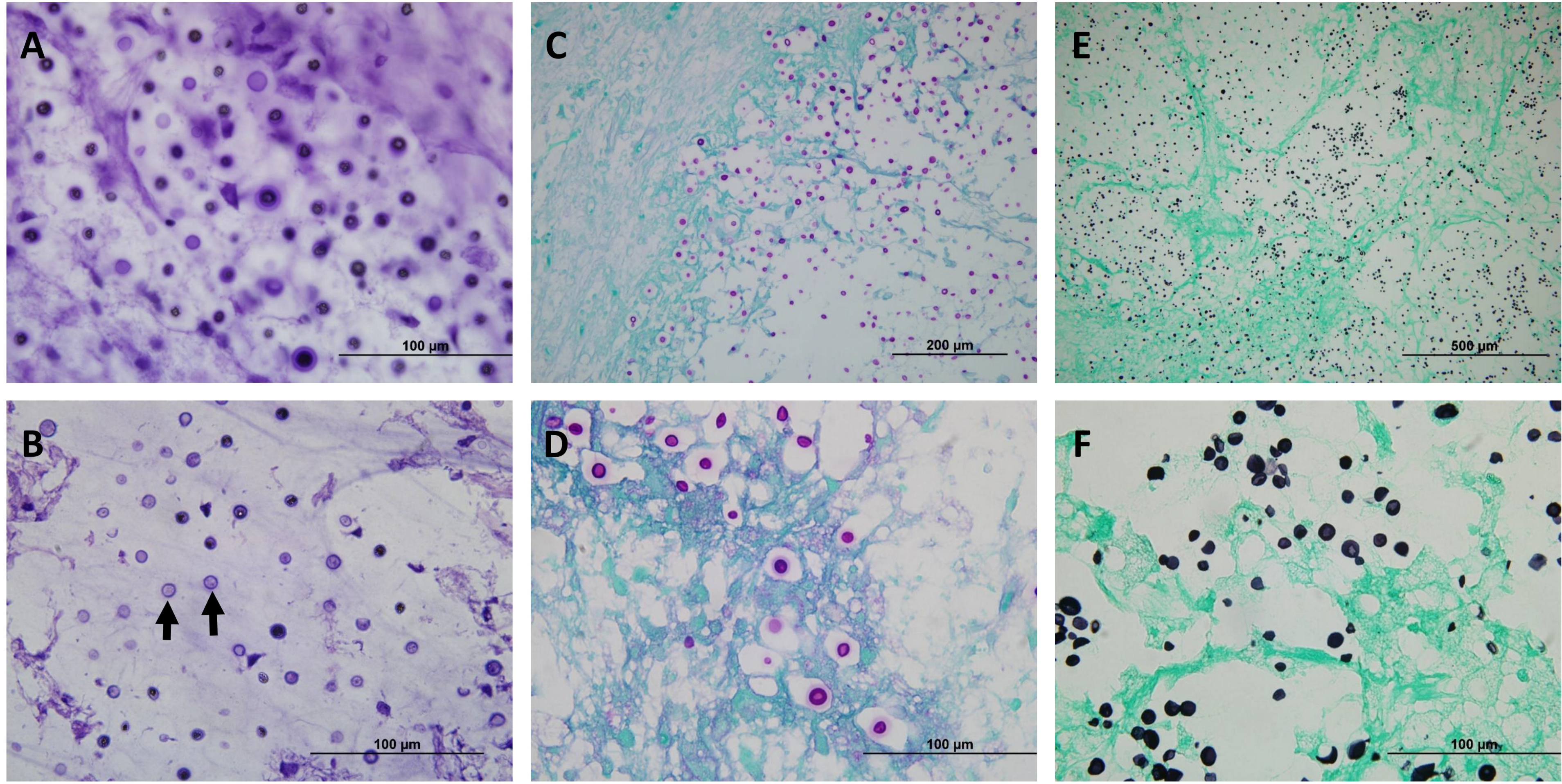
Cryptococcal organisms can be visualized within the brain parenchyma on both intraoperative and fixed tissues. A) Intraoperative touch preparation/smear depicting classic “soap bubble” lesions comprised of yeast cells within clear gelatinous capsules infiltrating the brain parenchyma. An absence of inflammatory cells was noted; the lack of an appreciable inflammatory response is characteristic of many cryptococcal infections. Scale bar represents 100 μm. B) Intraoperative frozen section demonstrated spherical-to-oval, variably sized, encapsulated yeast cells. The organisms were refractile and many had circumferential halos. Scale bar represents 100 μm. C) Low magnification image of Periodic Acid-Schiff (PAS) staining showed infiltration of brain parenchyma by positively-staining yeast cells, morphologically consistent with Cryptococcus. Scale bar represents 200 μm. D) Higher magnification of (C) highlighting the refractile appearance and the intensely stained mucopolysaccharide capsule of the yeast forms in tissue sections. Of note, mucicarmine and Alcian blue also showed positive staining in the capsule of cryptococcal organisms (not shown). Scale bar represents 100 μm. E) Low magnification image of Grocott-Gomori’s Methenamine Silver (GMS) staining showing a representative field with innumerable yeast cells, some with narrow-based budding. Scale bar represents 500 μm. F) Higher magnification of (E) illustrating the intense staining of organisms within the brain parenchyma (stained light green by the counter stain). Scale bar represents 100 μm.

A culture of the brain mass turned positive on day three of incubation. Matrix-assisted laser desorption/ionization time-of-flight (MALDI-TOF) analysis of the presumed fungal organism revealed a spectrum consistent with *C. gattii*. Immediately, a lumbar puncture was performed and cerebral spinal fluid (CSF) was sent for cryptococcal antigen testing. This returned positive (1:40), although culture of CSF was negative (Table 1). Fourth generation testing for HIV was negative, and a CD4^+^ T cell count obtained once the patient had completed the dexamethasone taper was normal at 871. Hemoglobin A1c was also within the normal range. Anti-GM-CSF testing was performed on the patient’s plasma and returned with high titers of these autoantibodies detected; in addition, elevated levels of immune system components interferon gamma (IFNγ) and interleukin-17A (IL-17A) were noted (Table 2).

**Table 2.** Screening of patient plasma for interferon gamma (INFγ), interleukin-17A (IL-17A), and anti-granulocyte-macrophage colony-stimulating factor (GM-CSF) autoantibodies. The positive controls are other patients with demonstrated titers and disease, while negative controls are healthy controls from a blood blank. The reported values are a measure of fluorescence intensity and are defined as relative light units (RLU).

| Sample | IFNγ | IL-17A | Anti-GM-CSF autoantibodies |
| --- | --- | --- | --- |
| Healthy Control (n=4) | 217 ± 263 RLU | 367 ± 461 RLU | 59 ± 46 RLU |
| Positive Control | 18085 RLU | 12850.5 RLU | 5397.5 RLU |
| Patient | 733.8 RLU | 731.3 RLU | 4711.3 RLU |

The patient improved clinically over the next four weeks of induction therapy, but then developed mild acute kidney injury (AKI) and cytopenia, prompting discontinuation of AmB and 5-FC, and transition to the renally-dosed equivalent of 800 mg fluconazole daily. Two days later, a repeat MRI of the brain was done to assess response to induction treatment and revealed an increase in size of the right parietal lesion and significant vasogenic edema (Figure 1B). AmB was then restarted in addition to fluconazole, and resection of the enlarging parietal lesion was performed. Pathology demonstrated a cryptococcoma with Grocott-Gomori’s methenamine silver (GMS) stain positive for fungal elements morphologically consistent with *Cryptococcus*. No culture was performed at the time of the second resection. The patient received an additional 48 hours of dexamethasone post-operatively, but as he was otherwise clinically stable no further steroid treatment was prescribed.

Two weeks later, after improvement in cytopenias, fluconazole was stopped and the patient was restarted on 5-FC with continuation of AmB. Over the following two weeks, the patient demonstrated continued slow improvement in symptoms, and brain MRI imaging showed stability of lesions (Figure 1C). Therefore, he was again transitioned to consolidation therapy with a renally equivalent dose of 800 mg fluconazole daily. After 12 additional weeks of high-dose fluconazole, maintenance therapy was started with fluconazole 400 mg daily. His neurologic symptoms gradually improved, with resolution of gait abnormalities and continued improvement in strength and coordination. As expected, AKI and cytopenias resolved with cessation of the AmB and 5-FC. A follow-up brain MRI completed after approximately 10 months of treatment revealed significant improvement (Figure 1D). Prolonged therapy with fluconazole is anticipated until complete resolution of the remaining intracranial lesions is achieved. A detailed timeline of the patient presentation, resections, and antifungal treatment is shown in Supplemental Figure 1.

### Phenotypic analysis places the clinical isolate in the *C. gattii* species complex

To confirm the MALDI-TOF results that characterized the clinical isolate as *C. gattii*, the isolate was subjected to a number of phenotypic tests. The isolate was grown on media to test for growth (YPD), melanin production (Niger Seed), urease production (Christensen’s Agar), and growth and color development on medium containing canavanine, glycine, and bromothymol blue (CGB). The isolate produced dark pigment on Niger Seed (melanin) and pink coloration on Christensen’s Agar (urease); both results are consistent with all pathogenic *Cryptococcus* species (Figure 3A). The isolate grew and produced blue pigment on CGB agar which is unique to *C. gattii* species (Figure 3A) (30). *Cryptococcus* produces a hydrophilic extracellular capsule as a means of evading the host immune system (31). To analyze capsule production, India ink staining and microscopic evaluation was performed after growth in CO_2_-independent medium for 72 hours at 37°C. Using the *C. deuterogattii* reference strain R265 for comparison, the patient isolate produced similar capsules (Figure 3B).

**Figure 3.**
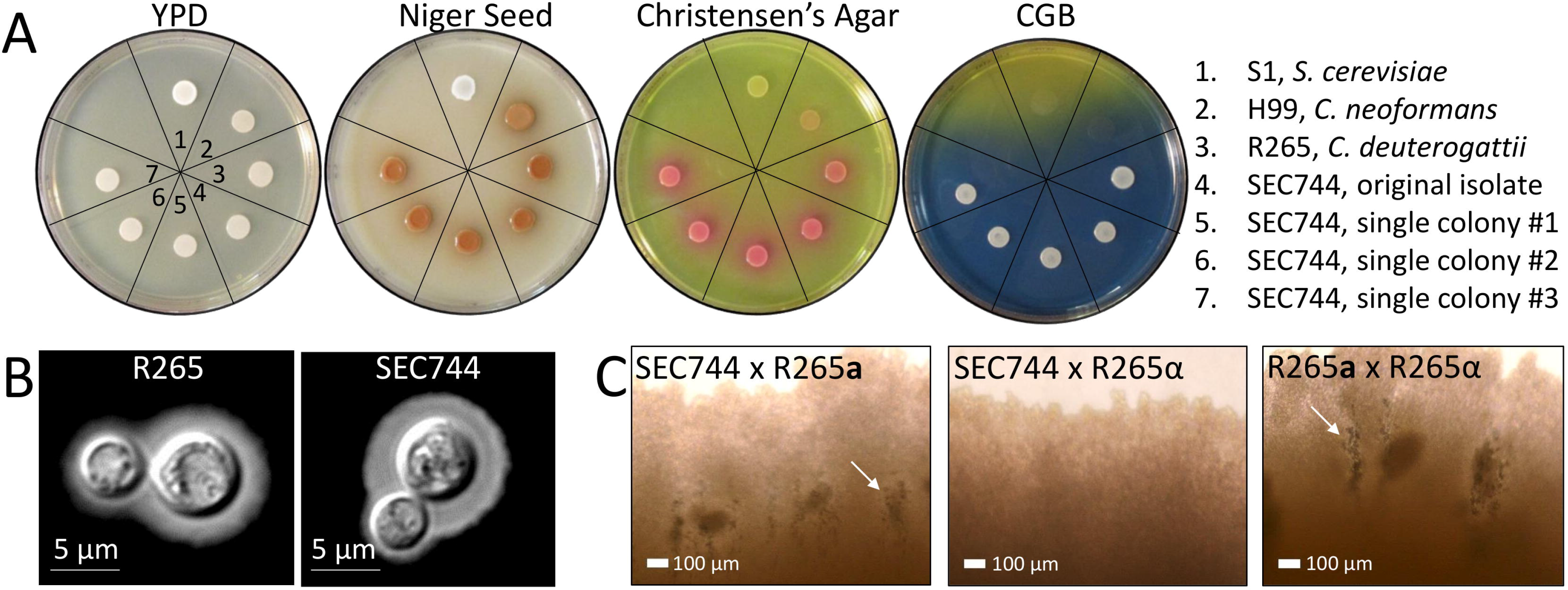
The clinical isolate displays phenotypes consistent with a *C. gattii MAT*α isolate. A) Phenotypic testing of the isolate on YPD, Niger Seed, Christensen’s agar, and canavanine, glycine, and bromothymol blue (CGB) media. 1 = S1 (*Saccharomyces cerevisiae*), 2 = H99 (*C. neoformans*), 3 = R265 (*C. deuterogattii*), 4 = Clinical isolate thick streak (SEC744), 5 = Clinical isolate single colony #1, 6 = Clinical isolate single colony #2, 7 = Clinical isolate single colony #3. B) Capsule staining with India ink. Scale bars represent 5 μm. C) Imaging of mating patches of the clinical isolate (SEC744) tested against R265 *MAT***a** and R265 *MAT*α. R265 *MAT***a** × α *MAT*α served as a positive control. Arrows indicate mating structures visible by light microscopy at 20X magnification. Scale bars represent 100 μm.

As previously mentioned, the majority of isolates characterized in *C. gattii* are of the *MAT*α mating type. To determine the mating type of the clinical isolate, it was co-cultured with α known *C. deuterogattii* VGIIa R265 *MAT***a** and *MAT*α isolates. The clinical isolate produced α mating structures (hyphae, basidia, and spores) only when paired with a *MAT***a** partner indicating the isolate is of the α mating-type (Figure 3C). No self-sporulation was observed when the patient isolate was incubated alone on mating media providing evidence that the isolate is not an **a-**α diploid or self-fertile (Supplemental Figure 2).

### Molecular analysis places the clinical isolate in the VGIIa major clade

MLST analysis was performed to assign the clinical isolate to a species within the *C. gattii* species complex. MLST loci were PCR amplified with primers designed to amplify each sequence, the PCR products were purified, analyzed by Sanger sequencing, and the sequences were assembled in Sequencher using forward and reverse strands to ensure complete double-stranded coverage. Phylogenetic analysis was performed using representative sequences from each of the *C. gatti* molecular types: *C. gattii* (VGI), *C. deuterogattii* (VGII), *C. bacillisporus* (VGIII), and *C. tetragattii* (VGIV). The gene sequences of the clinical isolate clustered closely with the *C. deuterogattii* representative sequence, supported with high bootstrap values (Figure 4A and Supplemental Figure 3). BLAST analysis of the contigs from each locus showed 100% sequence identity to the same loci in VGIIa sequences already deposited in NCBI. Allele designations were assigned for each of the loci, resulting in a clear pattern of the VGIIa genotype (Figure 4B). Taken together, the genotyping data assigns this clinical isolate into the VGIIa major lineage of *C. deuterogattii*. Fluorescence-activated cell sorting (FACS) was employed to determine that the isolate is haploid, containing a single copy of each chromosome (Figure 4C).

**Figure 4.**
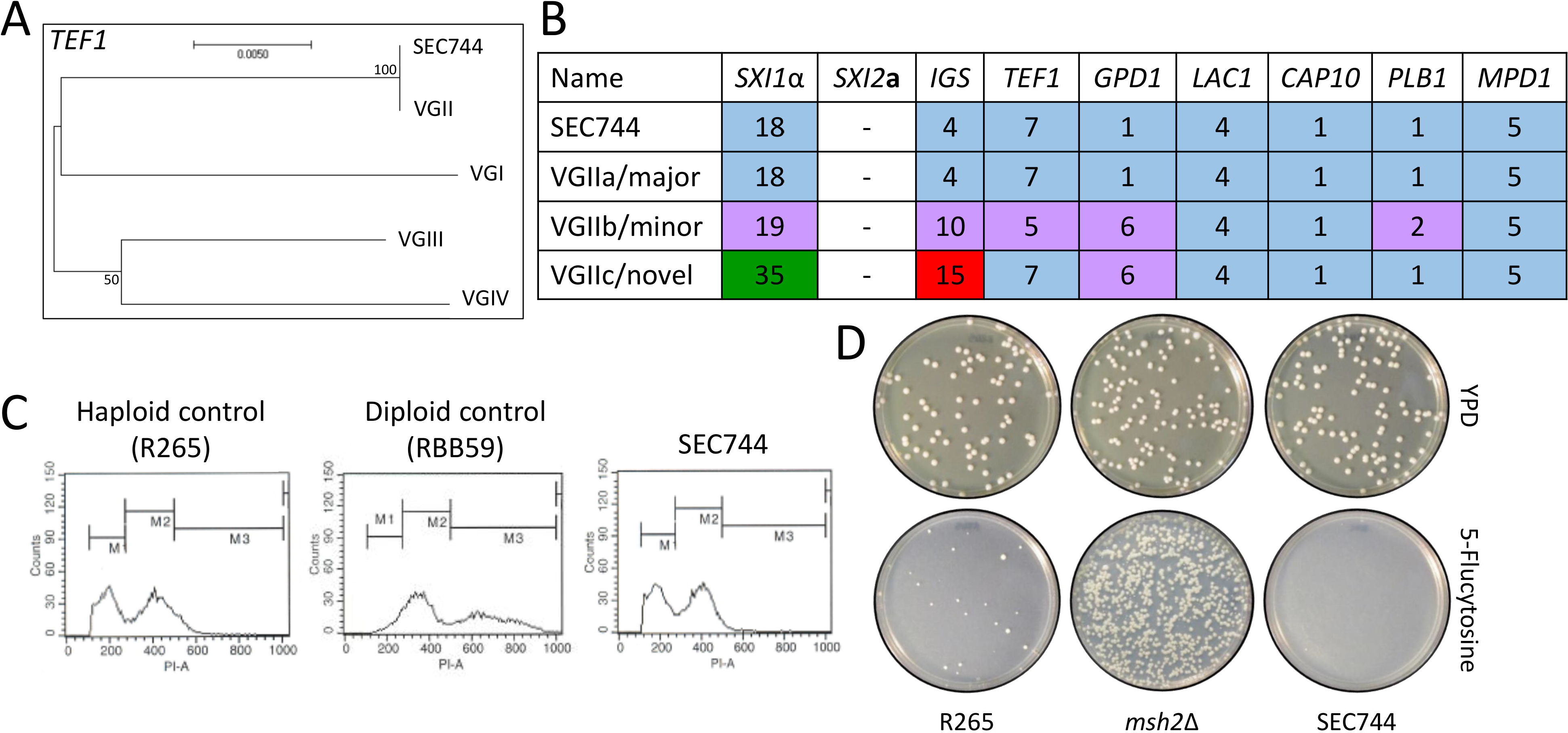
Genotyping places the clinical isolate in the *C. deuterogattii* VGIIa major lineage. A) Represenative MLST tree of *TEF1*. Scale bar represents the number of substitutions per site. B) Allele designations for the clinical isolate SEC744 and *C. deuterogattii* VGIIa, VGIIb, and VGIIc lineages. C) FACS analysis of *C. deuterogattii* R265 (haploid control), RBB59 (*C. gattii* diploid), and the clinical isolate SEC744. D) Testing for hypermutator phenotype in R265, *msh2*Δ, and the clinical isolate SEC744.

A lineage of *C. deuterogattii* VGIIa-like isolates was previously described that display a hypermutator phenotype resulting from a loss-of-function mutation in the *MSH2* gene, the MutS homolog required for mismatch DNA repair (32). Hypermutator strains display increased resistance to the fluorinated orotic acid derivative 5-fluoroorotic acid (5-FOA), the immunosuppressant FK506, and the clinically relevant antifungal drug 5-FC, suggesting a mode of resistance to antifungal therapy (32). To determine if the clinical isolate described here exhibits the same phenotype, the isolate was exposed to 5-FC and the ability of the strain to produce 5-FC resistant colonies was compared to wild-type and hypermutator control strains. Unlike the *de novo msh2*Δ mutant that produced a large number of 5-FC resistant colonies, the clinical isolate did not produce any colonies on medium supplemented with 5-FC (Figure 4D). This observation shows that the isolate displays a largely wild-type level of intrinsic mutation and is not a member of the hypermutator lineage.

### Clinical isolate shares its origin with *C. deuterogattii* outbreak strains

The isolate was subjected to whole genome sequencing and the resulting sequence was used in phylogenetic analysis with other previously described *C. deuterogattii* outbreak strains from the PNW (15). The clinical isolate clustered with ten highly clonal outbreak strains, including the VGIIa major reference strain R265 (Figure 5A). In accord with the phenotypic data, it did not fall within the hypermutator lineage that includes NIH444, CBS7750, and ICB107 (32). Alignment of the clinical isolate to the R265 reference strain revealed a total of 32 single-nucleotide polymorphisms (SNPs) that differ between the two strains. Of these SNPs, 22 were shared with at least one other strain in the outbreak group whereas ten SNPs were unique to the clinical isolate (Figure 5B). Of the ten private SNPs, seven were intergenic or intronic mutations and three were missense mutations that result in an amino acid change in the protein coding sequence. These nonsynonymous amino acid substitutions are all in uncharacterized genes and likely had no impact on the patient’s response to clinical treatment (Figure 5C).

**Figure 5.**
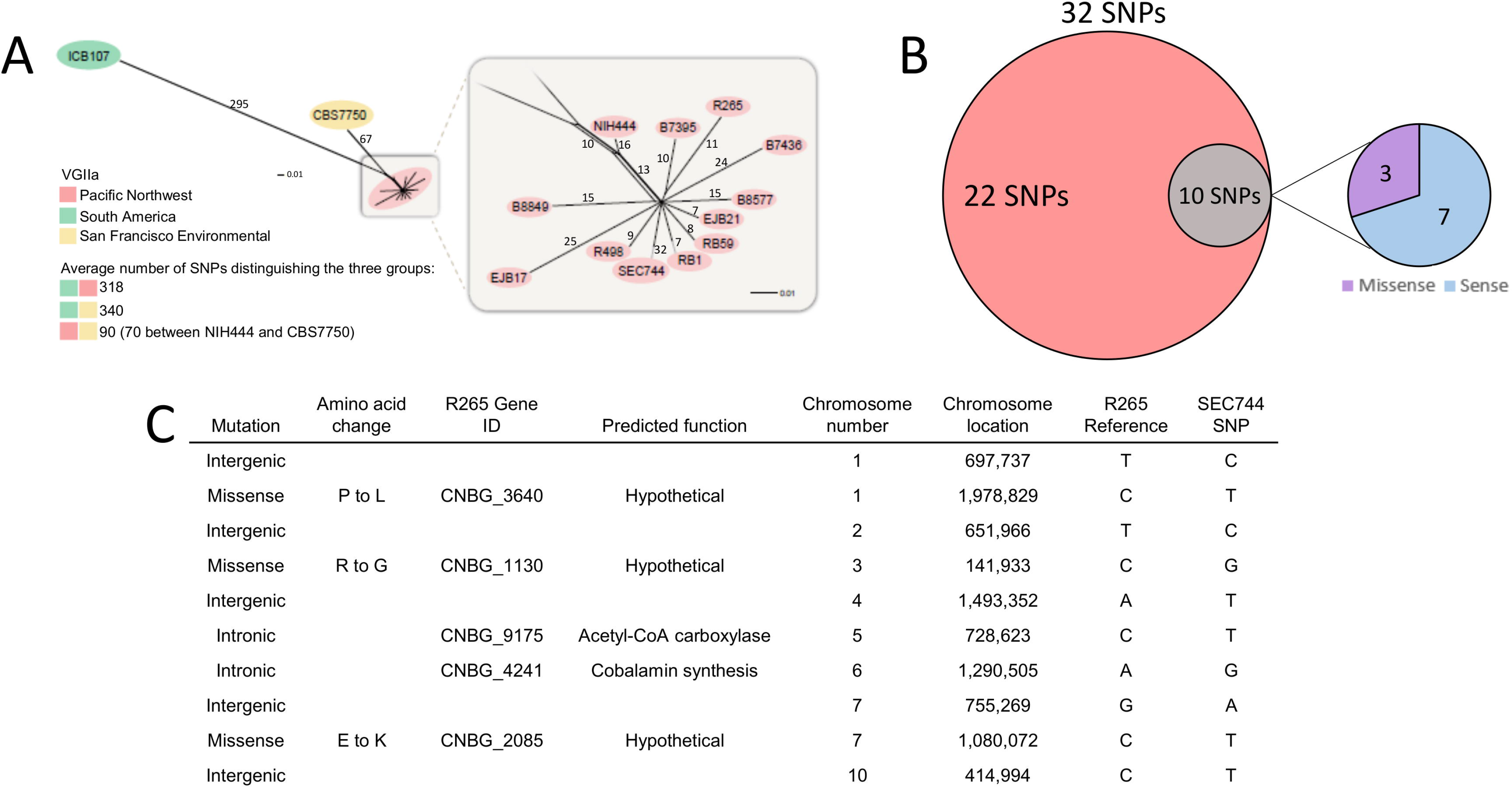
Phylogenetic analysis suggests the origin of the clinical isolate to be in the PNW. A) Phylogenetic tree depicting association of the clinical isolate SEC744 with other *C. deuterogattii* VGIIa outbreak strains from the PNW. Numbers on branches represent the number of SNPs distinguishing the groups. B) Venn diagram illustrating the SNPs found in the clinical isolate SEC744 compared to the outbreak VGIIa R265 reference strain. C) Table classifying the location and nature of the ten private SNPs in the clinical isolate SEC744.

## Discussion

In this study, we document the first case of a previously healthy individual becoming infected with *C. deuterogattii* after travel from the United States to the PNW outbreak region. Three largely clonal groups of isolates of *C. deuterogattii* have been identified circulating within the PNW: VGIIa/major which causes the majority of human and animal infections, VGIIb/minor which causes fewer infections, and VGIIc/novel, which so far has only been reported from Oregon, but not Seattle or Canada (21). More recent studies have suggested a South American origin of *C. gattii* where it is endemic, which makes the outbreak in the PNW exceptional given the temperate climate (33, 34). A 2018 study by Roe and colleagues estimated *C. gattii* was introduced to the PNW between 60 and 100 years ago, coinciding with the completion of the Panama Canal and the expansion of international shipping out of South America (35). However, *C. deuterogattii* VGIIb more closely aligns with isolates from Australia, suggesting multiple possible routes, sources, and introductions. A report from 2009 by Byrnes and colleagues documented a case of a travel-acquired *C. gattii* (VGI) infection (36), and since then three international travel-related *C. deuterogattii* (VGII) cases from individuals from Denmark, The Netherlands, Germany, and Japan have been reported (25–28). Overall, there is a noticeable increase in the number of travel-related cases being reported. Combined with the *C. deuterogattii* infection described here, this continues to bring an elevated level of attention to cryptococcal infections.

Following clinical presentation, a fungal mass was removed from the brain of the patient and subjected to phenotypic and genotypic analyses. Using microbiological techniques, the patient fungal isolate was identified as *C. deuterogattii* and detailed MLST and whole genome sequencing confirmed this finding. Further phylogenetic analysis shows the patient isolate clusters closely with previously described PNW outbreak strains (15). Of the strains identified as part of the PNW outbreak, three of the strains were characterized as being hypermutators due to their ability to produce drug resistant colonies at a high frequency. Following exposure to the clinically relevant 5-FC, the patient isolate did not produce a large number of drug resistant colonies suggesting it is not a member of this hypermutator lineage (32). Whole genome sequencing revealed 32 SNPs differentiating the patient isolate from the R265 reference strain. 22 of the SNPs were also found in at least one outbreak strain, but the clinical isolate contained 10 unique SNPs. These private SNPs do not appear to have influenced the pathogenicity of the infecting strain or treatment outcome as most are intergenic or resulted in amino acid changes in uncharacterized hypothetical proteins unlikely to play a significant role in disease (Figure 5).

When taken together, the clustering of the patient sample with the ten other highly clonal outbreak strains and the absence of a hypermutator phenotype provides strong evidence the origin of the patient isolate is within the PNW, and the patient acquired the infection during his visit to the area three years prior, which laid dormant until symptoms began following return to North Carolina. To our knowledge, this is the longest incubation period for a human *C. gattii* infection reported in the literature. However, there is currently limited evidence available to support an accurate estimate. Review of cases from the outbreak in the PNW suggests an incubation period of two to eleven months with a median of six to seven months (11). Although the patient also had travel history to a bird sanctuary in Costa Rica four months prior to the onset of symptoms, this seems less likely to be the source of infection as *C. deuterogattii* VGIIa is not known to be endemic in this area.

Cryptococcal meningitis in healthy individuals is somewhat rare (1 in 100,000), but the presentation of such cases suggests normal hosts could harbor rare primary immunodeficiences (37). Cryptococcal meningitis has been associated with a number of iatrogenic and inherited immunosuppressive conditions, including organ transplantation, hematopoetic malignancies, and steroid use. Furthermore, a number of defects in cell-mediated immunity have been identified as predisposing factors for cryptococcal meningitis. The most common immunodeficiency in non-HIV cryptococcal meningitis is idiopathic CD4^+^ lymphopenia, a disease caused by varying alleles, which is seen in over 25% of cases (37). Additionally, the presence of anti-GM-CSF and IFN-γ autoantibodies have also been associated with cryptococcal disease (10, 38). Monogenic mutations, such as those in the zinc transcription factor *GATA2*, the transcriptional activator *STAT3*, and the CD40 ligand *CD40LG*, are also associated with cryptococcosis in seronegative HIV patients (39–41). The patient described in this study did not have any documented iatrogenic causes of immunosuppression or history of immunodeficiency. Many hyper IgM syndromes often present earlier in life, in infancy or childhood, as increased susceptibility to sinopulmonary and opportunistic infections; this patient did not have such a history. Whole genome sequencing or exome sequencing of the patient could have determined if any of these underlying non-HIV related predisposing factors played a role in the patient’s acquisition of disease but were not performed in this study. Additionally, there is data to suggest that total and memory B-cell levels are lower in HIV-uninfected patients with *C. neoformans* infections (42). However, only limited flow cytometry measuring T-cell but not B-cell populations was analyzed in the patient.

Interestingly, the patient did have high titer anti-GM-CSF autoantibodies in the blood, which are known to cause acquired pulmonary alveolar proteinosis (PAP) through dysregulation of macrophage function, a cell type important for the control of cryptococcal infection. Specifically, these antibodies block STAT5 phosphorylation, which inhibits macrophage signaling (10, 29). Biologically inhibitory anti-GM-CSF autoantibodies have been associated with cryptococcal meningitis in some otherwise immunocompetent patients and in those with idiopathic CD4^+^ lymphopenia. However, it has yet to be confirmed if they are induced by an unknown trigger, preceding and therefore increasing susceptibility to infection, or if they are produced in response to infection. There is a notable absence of high titer anti-GM-CSF autoantibodies among the vast majority of healthy and immunodeficient control patients without cryptococcal disease reported thus far in the literature (29). Conversely, a recent study did find one healthy control who had anti-GM-CSF antibodies. His plasma inhibited STAT5 production, but to a much lesser degree than those with cryptococcal infection and autoantibodies (10). Because screening for anti-GM-CSF autoantibodies is not routinely performed in healthy individuals, for the patient described in this case it is not possible to know whether the autoantibodies predated or followed the acquisition of *C. deuterogattii*. The patient was screened for PAP in the clinical evaluation but a bronchoscopy was not performed as his symptoms did not warrant it (ie. no progressive dyspenea on exertion, chronic cough, weight loss, or fatigue). As of 15 months after the initial diagnosis he has not had any symptoms or CT chest imaging consistent with PAP.

Among HIV-uninfected patients, intracranial infection caused by *C. gattii* is associated with delayed response to therapy and higher incidence of neurosurgical intervention as compared to *C. neoformans* (43). In all patients, *C. gattii* infection is more likely to cause intracranial cryptococcomas, which often respond poorly to antifungal treatment. Guidelines recommend the same induction, consolidation, and suppressive treatment for infections due to both cryptococcal species, with extension of induction therapy to 6 weeks in those with cryptococcomas. In addition, surgery should be considered for lesions ≥ 3 cm with mass effect, those that fail to reduce in size after 4 weeks of therapy, or if there is compression of vital structures (2010 IDSA guidelines). Use of adjunctive corticosteroids in the setting of mass effect or vasogenic edema is supported by moderate evidence, but it is of limited (Level III) quality. The patient described here had initial clinical improvement in response to partial resection, steroids, and four weeks of induction therapy, but demonstrated radiographic deterioration with enlargement of the lesions and surrounding edema. This could have been due to completion of the steroid taper (last dose was nine days prior to MRI demonstrating increased lesions size), incomplete response to induction therapy especially in the setting of potential immunodeficiency, or an immune reconstitution inflammatory response-like syndrome that has been reported during treatment of *C. gattii* infection (44). While pathology did note fungal elements morphologically consistent with *Cryptococcus,* the GMS stain does not differentiate between live and dead organisms. Unfortunately, at the time of repeat resection, no culture of the lesion was done to determine if radiographic worsening was due to sterile inflammation or persistent infection. Since the second neurosurgical procedure, the patient has continued to recover clinically and has also demonstrated significant radiologic improvement. While he did receive two days of steroids post-operatively, it is unlikely that this would have resulted in the gradual, sustained improvement that ultimately followed. The slow progress suggests that although the second debulking surgery likely played a key role in his management, prolonged antifungal therapy was crucial to his treatment.

Disease caused by *C. gattii* can differ from that of *C. neoformans* based on host risks and clinical presentation. In this clinical case, a previously immunocompetent male presented with brain lesions, tested positive for anti-GM-CSF antibodies, and responded relatively well to treatment. Due to the extended dormancy stage of this infecting strain (~3 years) detailed travel history was key in implicating the cause and geographical source of the brain lesion. The need for detailed travel history with heightened clinical suspicion is critical as *C. gattii* often presents in different hosts than *C. neoformans*. This case also represents another example of anti-GM-CSF autoantibodies in a host concurrent with *C. gattii* infection, further strengthening the argument for anti-GM-CSF autoantibodies as a risk factor for *C. gattii* infection. Increased awareness will enhance the identification of diseases by treating physicians. In turn this will lead to more appropriate, precise, and timely treatments, and ultimately reduced morbidity and mortality.

## Materials and Methods

### Magnetic Resonance Imaging (MRI)

Imaging was performed using gadolinium contrast medium.

### Pathology

Intraoperative smear and frozen sectioning was performed at 60X magnification using hematoxyline and eosin. Periodic Acid-Schiff (PAS) staining and Grocott-Gomori’s Methenamine Silver (GMS) staining was performed at 20X and 60X magnification using formalin-fixed, paraffin-embedded tissue.

### Phenotypic tests

Strains were grown overnight in liquid YPD (1% yeast, 2% peptone, 2% dextrose), washed with sterile dH_2_O, and spotted onto solid YPD agar (1% yeast, 2% peptone, 2% dextrose, 1.5% agar), Niger Seed agar (7% niger seed, 0.1% glucose, 2% agar), Christensen’s Agar (0.1% gelatine peptone, 0.1% dextrose, 0.012 g phenol red, 0.2% KH_2_PO_4_, 1.2% agar), and canavanine, glycine, and bromothymol blue (CGB) agar (10% Solution A [1% glycine, 0.1% KH_2_PO_4_, 0.1% MgSO_4_, 1 mg Thiamine-HCl, 30 mg L-Canavanine sulfate], 2% Solution B [2% agar, 2% agar]). Plates were incubated at 30°C for two to three days.

### Capsule staining

The control strain R265 and the clinical isolate SEC744 were inoculated into liquid YPD (1% yeast, 2% peptone, 2% dextrose) and grown overnight at 30°C with rotation. Cultures were diluted 1:10 into 5 mL of CO_2_-independent medium and incubated for 3 days at 37°C with rotation. 500 μl of cells were pelleted and resuspended in 50 μl in sterile water. Following resuspension, 1.5 μ of cells were spotted onto a microscope slide along with 1 μl of India ink. Capsule staining was visualized using brightfield microscopy on a Ziess Axioscop 2 Plus microscope.

### Mating assay

Using sterile toothpicks the clinical isolate SEC744, R265 *MAT*α, and R265 *MAT***a** were patched together onto V8 pH=5 (5% V8 juice, 0.05% KH_2_PO_4_, 4% agar) as shown in Figure 2B. Mating plates were incubated in the dark at room temperature for 1 month and mating patches were observed under a Nikon Eclipse E400 microscope at 40X magnification.

### Phylogenetic analysis

Genes were amplified using primers listed in Supplemental Table 1 with typical PCR conditions. PCR products were purified and sent for Sanger sequencing (Genewiz, Research Triangle Park, NC). Consensus sequences were built using Sequencher with forward and reverse strands. Gene phylogenies were built using Mega7 Software with 100 bootstrap values using the Maximum Likelihood method based on the Tamura-Nei model (45). Representative consensus sequences for VGI, VGII, VGIII, and VGIV were acquired from GenBank entries.

### Multi-locus sequence typing (MLST)

To determine the molecular subtype of the clinical isolate SEC744, we aligned the consensus sequence from the phylogenetic analysis in Figure 4A and Supplemental Figure 3 to GenBank entries using Mega7 Software. If a 100% homologus alignment was found the allele was assigned with the already established allele designation numbers from Fraser *et al, Nature* 2005 (46). MLST gene sequences were submitted to GenBank and accession numbers will be assigned.

### Fluorescence-activated cell sorting (FACS)

Actively growing cells were scraped from a YPD agar plate (1% yeast, 2% peptone, 2% dextrose, 1.5% agar), washed with 1 mL PBS, and fixed overnight at 4°C in 1 mL 70% ethanol. Cells were then washed with 1 mL NS buffer (10 mM Tris-HCl pH 7.6, 250 mM sucrose, 1 mM EDTA pH 8.0, 1 mM MgCl_2_, 0.1 mM CaCl_2_, 0.1 mM ZnCl_2_, 0.4 mM phenylmethylsulfonyl fluoride, 7 mM β-mercaptoethanol) and then resuspended in 180 μL NS buffer with 5 μl propidium iodide (0.5 mg/mL) and 20 μl RNase (10 mg/mL). Cells were stained overnight at 4°C. Prior to analysis on the flow cytometer, 50 μl mM Tris-HCl pH8.0 in a 5 mL Falcon tube. Flow cytometry was performed on 10,000 cells and analyzed on the FL1 channel on a Becton-Dickinson FACScan.

### Hypermutator assay

Single colonies were inoculated into a 5 mL overnight YPD liquid culture (1% yeast, 2% peptone, 2% dextrose). Cells were pelleted, washed with 5 mL dH_2_O, and resuspended in 5 mL dH_2_O. 100 μl of a 10^−5^ dilution was plated on YPD agar (1% yeast, 2% peptone, 2% dextrose, 1.5% agar) to check cell viability and 100 μl of undiluted culture was plated on YNB+5-FC (0.67% yeast nitrogen base, 2% glucose, 2% agar, 100 μg 5-FC).

### DNA extraction and genome sequencing

Genomic DNA was extracted using cetyltrimethylammonium bromide (CTAB) buffer. A 25 mL culture of SEC744 was pelleted and lyophilized before pulverizing with glass beads. The dried cell debris was resuspended in 10 mL CTAB buffer and heated at 65°C for 10 minutes. An equal volume of chlorofom was added to the mixture and the solution was gently mixed. Following centrifugation, the aqueous phase was transferred to a new tube and DNA was precipitated using isopropanol. The DNA was pelleted, resuspended in dH_2_O and treated with 20 μg/mL of RNase A. Whole genome sequencing was performed at the Duke University Sequencing and Genomic Technologies Shared Resource (Durham, North Carolina). Libraries were prepared using the Kapa Hyper Prep Kit, and the libraries were run on an Illumina Hi-Seq 4000 Instrument to generate 150 base pair paired-end reads.

### SNP calling and phylogenetic networks analysis

The phylogenetic relationship of the clinical isolate SEC744 with other strains of the *C. deuterogattii* VGIIa subpopulation was inferred from a matrix containing the SNP calls for the selected set of samples. The genomic paired-end reads were mapped to the *C. deuterogattii* R265 reference genome (GenBank accession: GCA_002954075.1) (47) using BWA-MEM short read aligner v.0.7.17-r1188 (48) with default settings. SNP discovery, variant evaluation, and further refinements were carried out with the Genome Analysis Toolkit (GATK) best practices pipeline (49) v.4.0.1.2, including Picard tools to convert SAM to sorted BAM files, fix read groups (module: AddOrReplaceReadGroups; SORT_ORDER=coordinate), and mark duplicates. Variant sites were identified with the HaplotypeCaller from GATK using the haploid mode setting and subsequently filtered using the VariantFiltration module with the following criteria: QD < 2.0 || FS > 60.0 || MQ < 40.0 || MQRankSum < −12.5 || ReadPosRankSum > −8.0 || SOR > 4.0. Next, variants were parsed with VCFtools v0.1.16 (50) to include only high-quality SNPs, here defined as SNPs with mean depth >10 (--minDP 10) and minimum quality of 30 (-- minQ 30). The option --max-missing 1 was used to exclude sites with missing data in at least one sample. The resulting VCF files were parsed with a custom Perl script into a concatenated FASTA consisting of 526 sites and then imported to SplitTree v.4.14.2 (51) to construct an unrooted split-network using the Neighbor-net method. The potential mutational effect of the 10 private variants found in the clinical isolate SEC744 was manually inspected by substituting the corresponding bases in the annotated genes of the *C. deuterogatii* R265 with these variants.

### Nucleotide sequencing accession number

The genomic reads generated here were deposited in the Sequence Read Archive under project accession number PRJNA493061.

### Anti-granulocyte-macrophage colony-stimulating factor (GM-CSF) autoantibody testing

Anti-GM-CSF autoantibody testing was performed at the National Institute of Allergy and Infectious Diseases in Bethesda, Maryland. Briefly, the ability of the antibody to directly bind to interferon gamma (INFγ), interleukin-17A (IL-17A), or anti-GM-CSF autoantibody was assessed and measured as a function of fluorescence intensity (52).

## Declaration of interests

The authors declare no competing interests.

## Acknowledgements

This work was supported by NIH/NIAID R01 grants AI50113-14 and AI112595-04 and NIH/NIAID R37 grant AI39115-21. We thank Sheng Sun, Shelby Priest, and Cecelia Wall for helpful comments on the manuscript.

---

**Supplemental Figure 1. Detailed timeline of the patient’s clinical course.**

**Supplemental Figure 2. Clinical isolate does not sporulate in solo culture.**

Clinical isolate SEC744, R265 *MAT*α, and R265 *MAT***a** were patched individually onto mating media (V8 pH=5). No hyphae or spore formation were observed.

**Supplemental Figure 3. Phylogenetic analysis of six loci in the clinical isolate cluster with *C. deuterogattii*.**

MLST trees of *IGS, LAC1, GPD1, CAP10, PLB1,* and *MPD1* show that these gene sequences are all most similar to the same loci in *C. deuterogattii* compared to *C. gattii* (VGI), *C. deuterogattii* (VGII), *C. bacillisporus* (VGIII), and *C. tetragattii* (VGIV). All results are supported by strong boostrap values. Scale bar represents the number of substitutions per site.

**Supplemental Table 1. Primers used in this study.**

