## Supplementary figures and images for "*Cryptococcus deuterogattii* VGIIa infection associated with travel to the Pacific Northwest outbreak region in an anti-GM-CSF autoantibody positive patient in the United States"

### Supplemental Figure 1

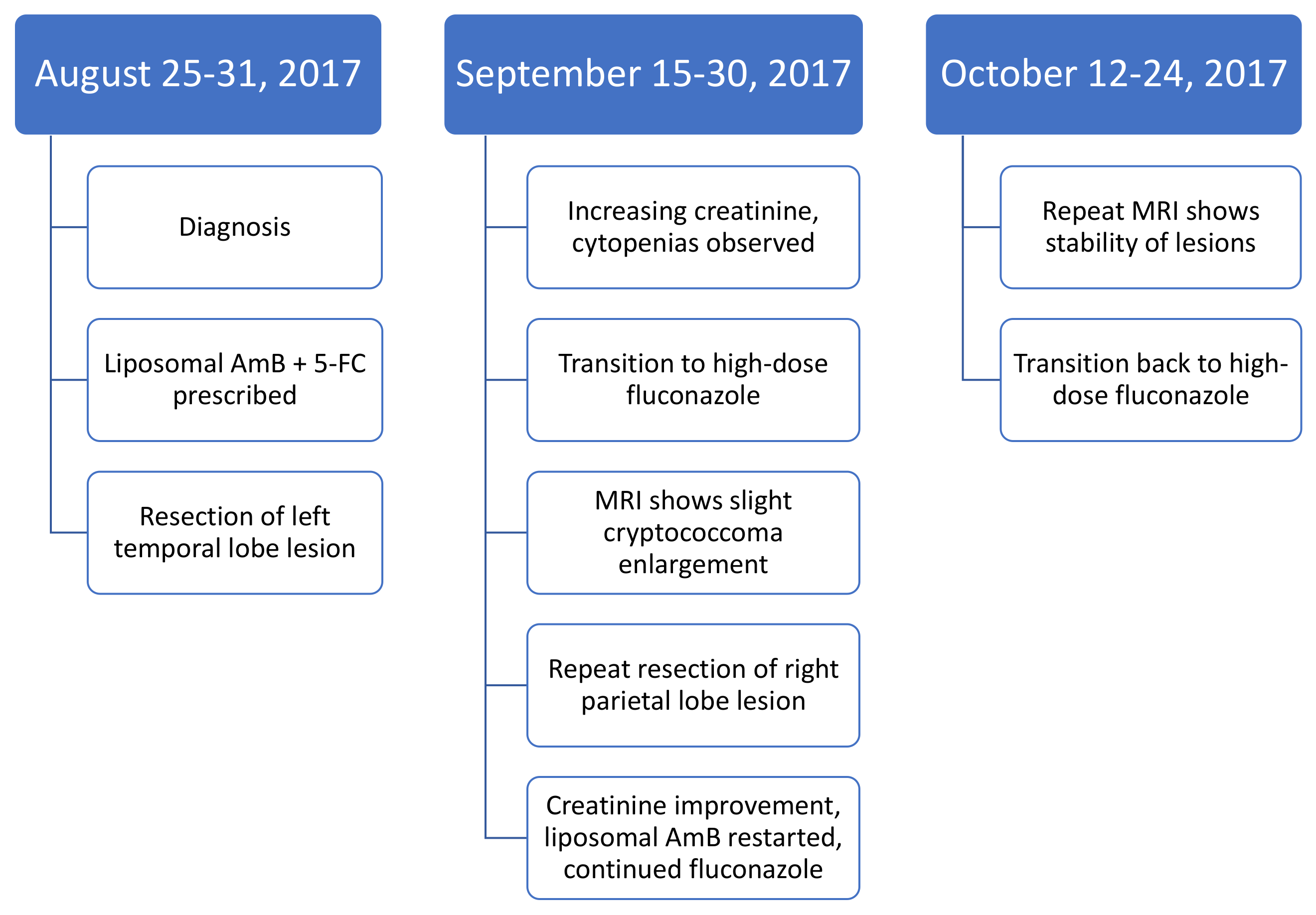

### Supplemental Figure 2

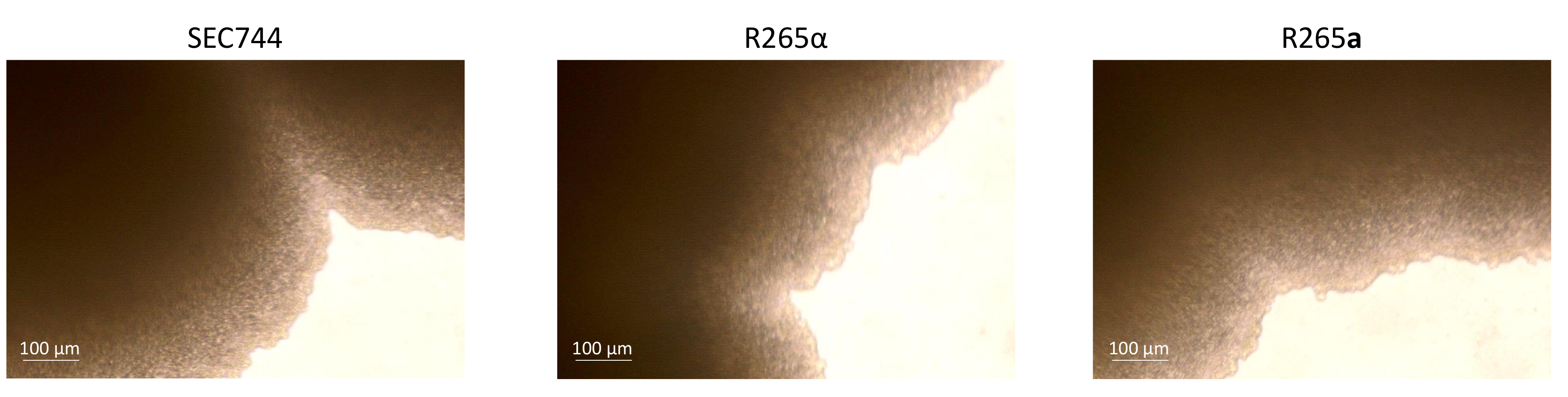

### Supplemental Figure 3

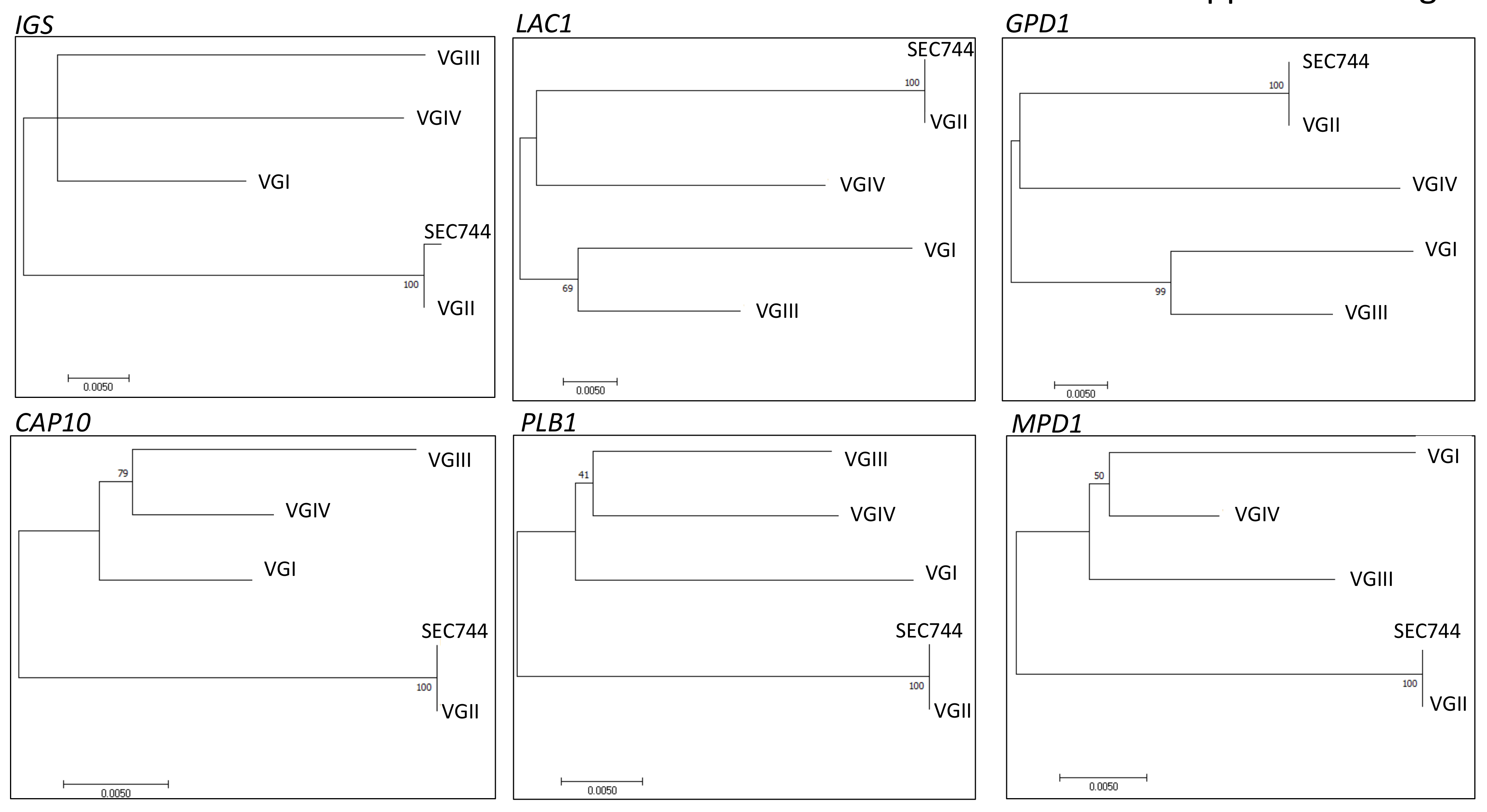
