## Supplemental Table 1 for "*Cryptococcus deuterogattii* VGIIa infection associated with travel to the Pacific Northwest outbreak region in an anti-GM-CSF autoantibody positive patient in the United States"

Supplemental Table 1. Primers used in this study

| Primer name | Sequence | Gene |
| --- | --- | --- |
| JOHE14115 | AGGGTACGTTTGAGGCCAGTT | *SXI1*α 5’ MLST primer |
| JOHE14116 | GAAAGCGTTGGCAAGGAATGA | *SXI1*α 3’ MLST primer |
| JOHE10453 | TGATCGCACGAGCCAAATCCC | *SXI2***a** 5’ MLST primer |
| JOHE10454 | GGCTTCCTGACAACACTTCTA | *SXI2***a** 3’ MLST primer |
| JOHE14408 | ATCCTTTGCAGACGACTTGA | *IGS* 5’ MLST primer |
| JOHE14409 | GTGATCAGTGCATTGCATGA | *IGS* 3’ MLST primer |
| JOHE14976 | GCACGCTCTTCTCGCCTTCAC | *TEF1* 5’ MLST primer |
| JOHE14977 | GTAGTCGGCGTAGGTCTCAAC | *TEF1* 3’ MLST primer |
| JOHE14968 | CCACCGAACCCTTCTAGGATA | *GPD1* 5’ MLST primer |
| JOHE14969 | CTTCTTGGCACCTCCCTTGAG | *GPD1* 3’ MLST primer |
| JOHE14970 | AACATGTTCCCTGGGCCTGTG | *LAC1* 5’ MLST primer |
| JOHE14971 | ATGAGAATTGAATCGCCTTGT | *LAC1* 3’ MLST primer |
| JOHE14386 | CCGGAACTGACCACTTCATC | *CAP10* 5’ MLST primer |
| JOHE14387 | GCCCACTCAAGACACAACCT | *CAP10* 3’ MLST primer |
| JOHE14974 | CTCTCATTGTTCGCCGCTACT | *PLB1* 5’ MLST primer |
| JOHE14975 | GGAAGCCGAGGTCTGATTTGG | *PLB1* 3’ MLST primer |
| JOHE14972 | TGCCCTGGATCCTAATGCTCT | *MPD1* 5’ MLST primer |
| JOHE14973 | ACCCAGACTGCCGCTGTCGTC | *MPD1* 3’ MLST primer |
